# Architecture and Energy Transfer of the Bacterial Photosynthetic Unit

**DOI:** 10.64898/2026.09.23.753705

**Authors:** Peng Wang, Ze-Kun Liu, Fan Xu, Jing-Li Lv, Meng-Qi Wang, Jian-Xun Li, Jiaxing Han, Feifei Li, Yingyue Zhang, Jun Gao, Yu-Zhong Zhang, Lu-Ning Liu

## Abstract

In phototrophic organisms, pigment-protein membrane complexes are densely packed to form photosynthetic units (PSUs) that capture solar energy and convert it into chemical energy. Although the structures of many individual photosynthetic complexes have been resolved, how they are arranged and interact with others within photosynthetic membranes to enable efficient excitation energy transfer (EET) remains poorly understood. Here, we report cryo-electron microscopy structures of PSU supercomplex assemblies from the phototrophic α-proteobacterium *Rhodovulum viride*, including an RC–LH1 core associated with one or two peripheral LH2 complexes and a curved LH2 tetramer. These membrane-derived assemblies define the relative positions and orientations of neighboring photosynthetic complexes and place their pigment arrays in proximity across antenna– antenna and antenna–core interfaces. Structure-based simulations identify potential EET pathways within the PSU assemblies and reveal rapid energy transfer across both LH2–LH2 and LH2–LH1 interfaces. Collectively, these findings provide insights into the assembly and structural modularity of bacterial PSUs and elucidate how the lateral organization of membrane protein complexes facilitates efficient energy transfer. This work extends structural studies of bacterial photosynthesis from individual complexes to their native higher-order assembly, providing a framework for understanding how photosynthetic supercomplex organization shapes energy migration and for guiding the design of artificial photosynthesis.

## Introduction

Photosynthetic organisms assemble light-harvesting antennae and reaction centers (RCs) into supercomplexes that couple photon absorption to charge separation, sustaining most life on Earth (1-3). Resolving the native architecture and protein-protein interactions of these photosynthetic supercomplexes is essential for understanding how the photosynthetic apparatus is generated and maintained to enable efficient excitation energy transfer (EET) and electron transport.

In the photosynthetic unit (PSU) of purple phototrophic bacteria, light-driven charge separation is initiated in anoxygenic Type II RCs, which are surrounded by light-harvesting complex 1 (LH1) to form RC–LH1 core complexes (1, 4-7). LH1 captures solar energy either directly or indirectly from peripheral antenna systems, most notably light-harvesting complex 2 (LH2), and funnels it to the RC, where sequential electron transfer generates a proton motive force to drive ATP synthesis (8-11). Although numerous high-resolution structures of isolated RC–LH1 and LH2 complexes have been determined individually (12-20), how these complexes are organized and interact within native PSUs to support rapid excitation energy transfer remains poorly understood.

Atomic force microscopy (AFM) has revealed lateral arrangements of LH2 antennae and RC– LH1 core complexes in native photosynthetic membranes, but their molecular interfaces remain incompletely defined (21-27). Recently, cryo-EM and ultrafast spectroscopy of reconstituted LH2/LH3 pairs in nanodiscs linked intercomplex pigment spacing to EET (28). In addition, an LH1– LH2 complex from *Halorhodospira halophila* shows an unusual concentric architecture in which an 18-subunit LH1 ring surrounds a nine-subunit LH2 ring without an RC (29). Despite these studies, how peripheral LH2 antennae associate with an intact RC–LH1 core in the bacterial PSU and how the supercomplex organization connects their pigment networks remain unclear.

Here, we report cryo-electron microscopy (cryo-EM) structures of the RC–LH1–LH2 supercomplexes and LH2 oligomeric assemblies isolated from an anoxygenic purple non-sulfur phototroph *Rhodovulum viride* (30). These membrane-derived structures reveal the molecular architecture of the photosynthetic assemblies and the physical associations of peripheral LH2 and the RC–LH1 core. The pigment arrangements further enable calculations of energy-transfer pathways within the photosynthetic assemblies by employing computational simulations. Collectively, these findings establish a structural framework for understanding the native organization and energy-transfer pathways of the bacterial PSU.

## Results and Discussion

### Identification of the physical association between RC–LH1 and LH2

Members of the genus *Rhodovulum* are representative anoxygenic purple non-sulfur phototrophs with several distinctive features (30). Their RC–LH1 complexes contain a cytochrome (Cyt) *c* subunit carrying three heme cofactors rather than the canonical four-heme configuration found in most purple bacteria, and they are capable of phototrophic growth under aerobic conditions (10, 20, 31). *Rdv. viride*, a model species within this genus, is further distinguished by its characteristic green pigmentation, offering an attractive system for investigating the adaptive mechanisms of bacterial photosynthetic machinery (32-34).

To investigate the structures of the photosynthetic complexes in *Rdv. viride*, photosynthetic membrane complexes were isolated from phototrophically grown cells by sucrose density-gradient ultracentrifugation (Fig. S1A). Two distinct green bands were reproducibly observed. The upper band showed LH2-specific absorption maxima at 800 and 850 nm, whereas the lower band exhibited a dominant peak at 880 nm with weaker 800- and 850-nm features, indicating predominantly RC–LH1 cores with a minor LH2 component that was barely detectable by SDS–PAGE (Figs. S1B, S1C).

We next performed single-particle cryo-EM analysis on both isolated LH2 and RC–LH1-enriched fractions. Reconstructions of isolated RC–LH1 and LH2 complexes were obtained at overall resolutions of 2.58 Å and 2.87 Å, respectively (Figs. 1, S2 and S3). Further 3D classification of the RC–LH1 dataset resolved two distinct supercomplex assemblies, one RC–LH1 core associated with one LH2 ring (RC–LH1–LH2) or two LH2 rings (RC–LH1–2LH2), at 5.68 and 6.77 Å, respectively, supporting the physical association of peripheral LH2 with RC–LH1 cores. An LH2 tetramer was also reconstructed at 7.72 Å (Fig. S2).

**Fig. 1.**
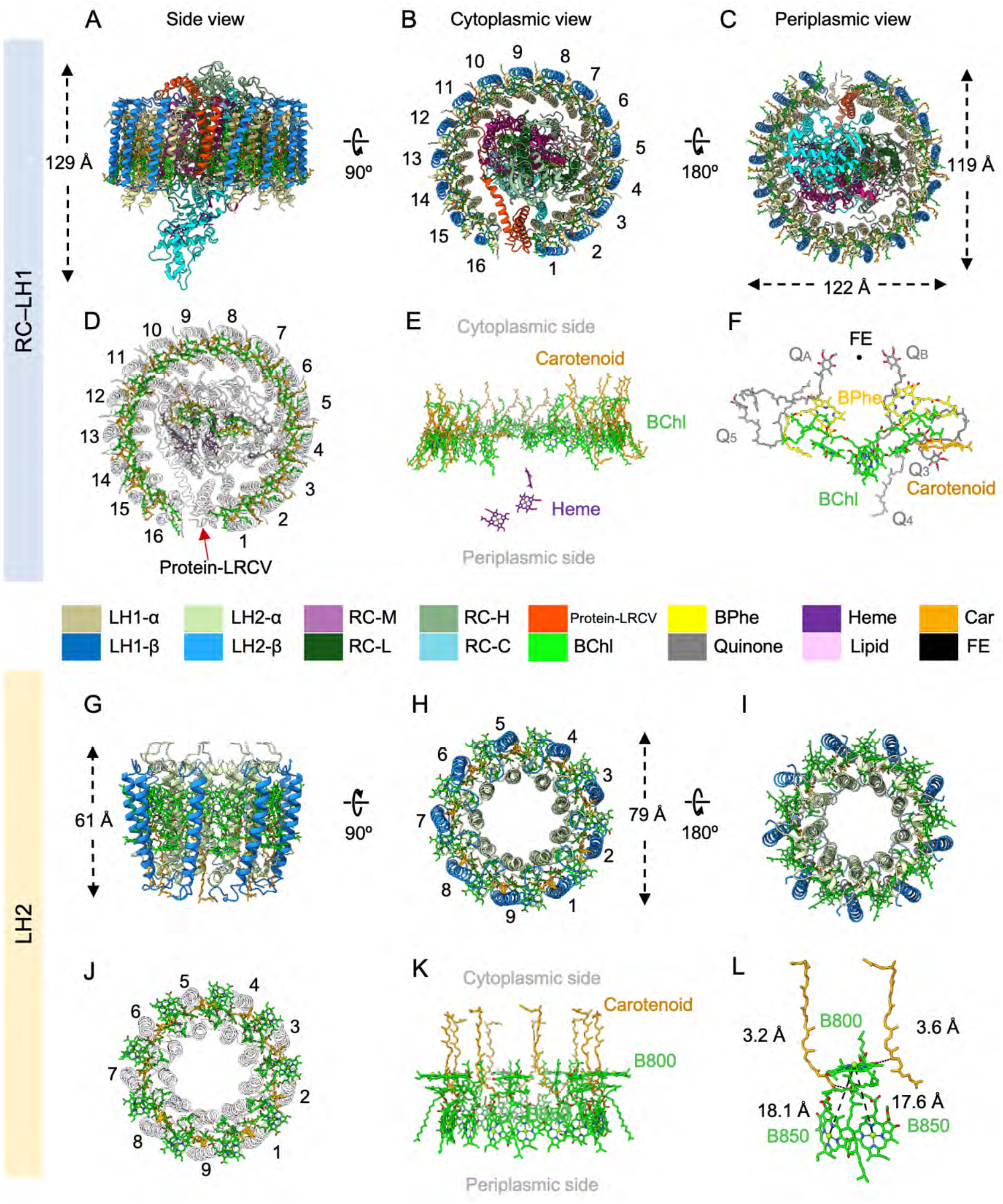
Cryo-EM structures of the RC–LH1 and LH2 complexes from *Rdv. viride*. (A to C) Structural model of the RC–LH1 complex in three different views, colored as indicated in the key at the bottom of the figure. (A) Side view of the RC–LH1 complex in the membrane plane. (B) Cytoplasmic view of the RC–LH1 complex. (C) Periplasmic view of the RC–LH1 complex. (D) Pigments and cofactors viewed from the cytoplasmic side in the RC–LH1 complex. (E) Side view of the arrangement of pigments and cofactors in the membrane plane in the RC–LH1 complex. (F) Arrangement of pigments, quinones, and lipids associated with the reaction center (RC). (G to I) Structural model of the LH2 complex in three different views, colored as indicated in the key at the top of the figure. (G) Side view of the LH2 complex in the membrane plane. (H) Periplasmic view of the LH2 complex. (I) Cytoplasmic view of the LH2 complex. (J) Pigments and cofactors viewed from the cytoplasmic side in LH2. (K) Side view of the arrangement of pigments and cofactors in the membrane plane in LH2. (L) Mg–Mg distances between B850 and B800 BChls, and distances between carotenoids and their adjacent BChls.

### Structures of individual RC–LH1 core complex and the LH2 complex

High-quality density maps of the RC–LH1 complex and LH2 enabled confident modeling of nearly all protein side chains, pigments, lipids and cofactors, providing complete atomic models of both photosynthetic modules (Figs. S4 and S5, Table S1).

The RC–LH1 complex from *Rdv. viride* features a central RC encircled by an incomplete LH1 ring (Figs. 1A-1C). The RC contains the conserved L, M, H and Cyt *c* subunits, together with four bacteriochlorophyll (BChl) *a* molecules, two bacteriopheophytins, three c-type hemes, five ubiquinone-10 molecules, one carotenoid, six lipid molecules and a non-heme iron (Figs. 1A-1F). The slightly elliptical open LH1 ring comprises sixteen αβ-heterodimers, with overall dimensions of approximately 122 × 119 Å in the membrane plane and 129 Å in height (Figs. 1A, 1C). Each αβ-heterodimer coordinates two BChl *a* molecules, while a total of thirty carotenoids occupy the spaces between adjacent αβ pairs. Together, these pigments form a continuous peripheral layer surrounding the RC (Fig. 1D).

The overall RC–LH1 architecture closely resembles that of the *Rdv. sulfidophilum* RC–LH1 complex (20), indicating a high degree of structural conservation within the genus (Fig. S6). Two notable features distinguish these *Rhodovulum* cores from many other RC–LH1 complexes. First, a unique transmembrane protein with three α-helices termed “Protein-LRCV” occupies the opening in the LH1 ring, interrupting the LH1 array and stabilizing the opening through interactions with adjacent LH1 and RC subunits, similar to its counterpart in *Rdv. sulfidophilum* (20) (Figs. 1D and S7). Second, the RC contains a three-heme Cyt *c* subunit rather than the four-heme configuration found in most purple phototrophic bacteria (Figs. 1E and S8).

The *Rdv. viride* LH2 adopts an architecture similar to that of *Rba. sphaeroides* (Fig. S9) (15). The LH2 complex forms a cylindrical nonamer approximately 79 Å in diameter and 61 Å in height (Figs. 1G-1I). Nine αβ heterodimers assemble into a closed ring surrounding a central cavity, and each heterodimer coordinates three BChl *a* molecules and one carotenoid, generating the characteristic B800-B850 antenna system (Figs. 1J-1L).

### Neurosporene defines the characteristic pigment composition of *Rdv. viride*

Green pigmentation is a defining feature of *Rdv. viride* (Fig. S1A), a phenotype shared with *Rba. viridis* but distinct from the yellow-brown coloration of most purple phototrophic bacteria (5, 32, 33). Previous biochemical analysis indicates that neurosporene, a nonaene carotenoid with a shorter conjugated π-electron system, is the predominant carotenoid in *Rdv. viride* (32, 33, 35). Early termination of the phytoene desaturase (CrtI) reaction limits desaturation, shifting carotenoid absorption and the macroscopic color (35).

Cryo-EM maps revealed well-defined densities for neurosporenes (Figs. S4 and S5). In RC– LH1, thirty molecules modeled as neurosporene locate between neighboring LH1 αβ-heterodimers and form a continuous ring surrounding the BChl *a* array (Fig. 1E). In LH2, each αβ-heterodimer accommodates one neurosporene located between the two transmembrane helices in close proximity to both B800 and B850 BChls (Figs. 1K, 1L). These contacts suggest the conserved light-harvesting and photoprotective roles of carotenoids in antennae of purple photosynthetic bacteria (36, 37).

### Architecture of the RC–LH1–LH2 photosynthetic unit

Intriguingly, we identified the co-purification of LH2 with RC–LH1 complexes, suggesting their stable association that persists even after membrane solubilization. Three-dimensional classification resolved multiple assemblies in addition to RC–LH1 only (Figs. 2, 3 and S2): (i) an RC–LH1 core associated with one LH2 (RC–LH1–LH2), (ii) an extended assembly containing two LH2 complexes (RC–LH1–2LH2), and (iii) an LH2 tetramer. Although the map resolution is insufficient for *de novo* atomic model building, rigid-body fitting of the obtained high-resolution RC–LH1 and LH2 structures defined the relative positions and orientations of the component complexes within the PSU (Figs. 1 and S10).

**Fig. 2.**
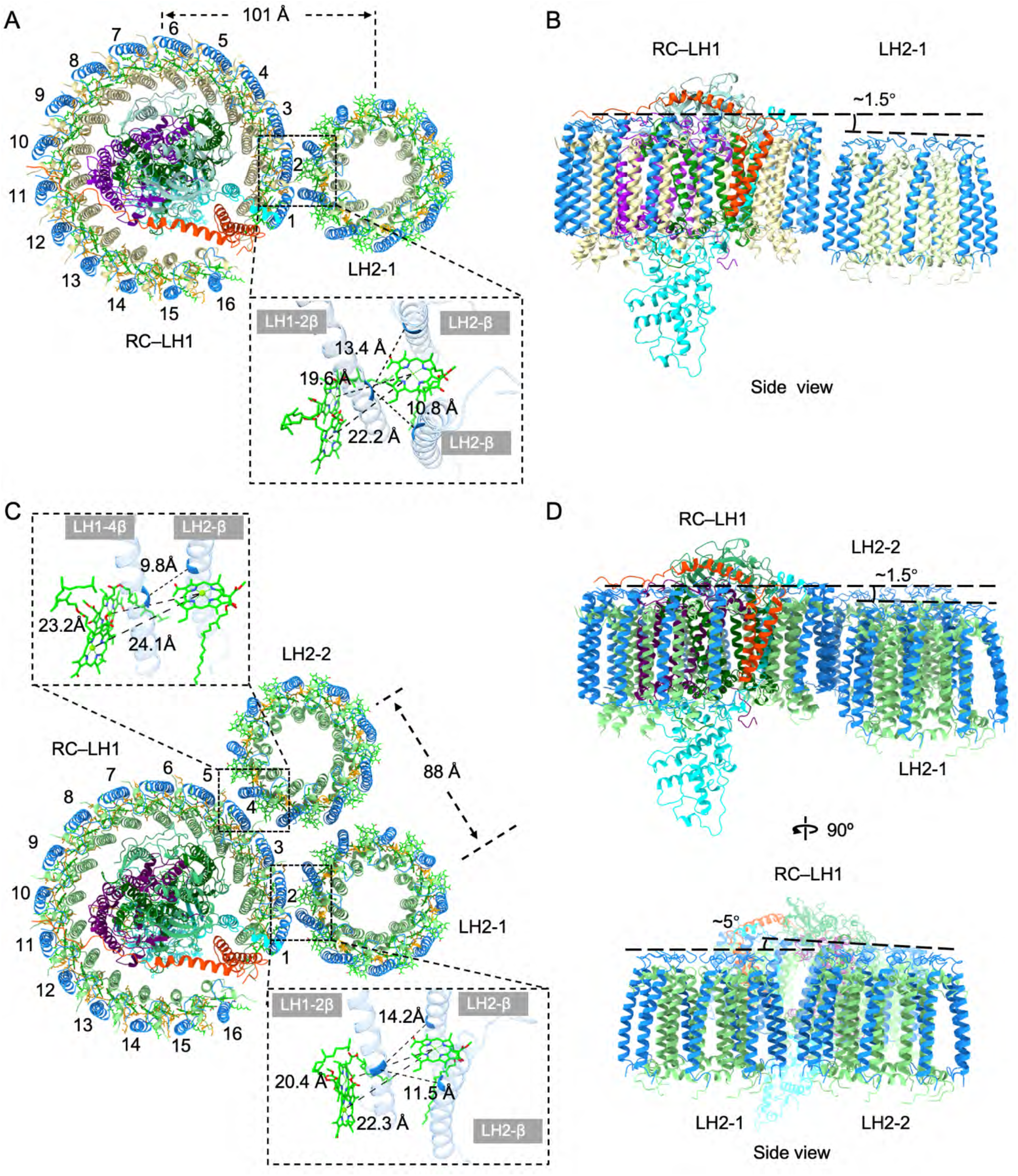
Structural organization of the RC–LH1–LH2 and RC–LH1–2LH2 assemblies. (A) Cytoplasmic view of the RC–LH1–LH2 assembly. The interaction interface between RC–LH1 and LH2 is shown in the zoomed-in view. (B) Side view of the RC–LH1–LH2 assembly, showing a relative tilt angle of approximately 1.5° between the two complexes. (C) Cytoplasmic view of the RC–LH1–2LH2 assembly. The second LH2 complex is incorporated without large-scale rearrangements of the RC–LH1 core and neighboring LH2. The interaction interfaces between RC– LH1 and LH2-1 or LH2-2 are shown in the zoomed-in views. (D) Side views of the RC–LH1–2LH2 assembly, showing a relative rotational offset of approximately 5° between the two LH2 complexes.

**Fig. 3.**
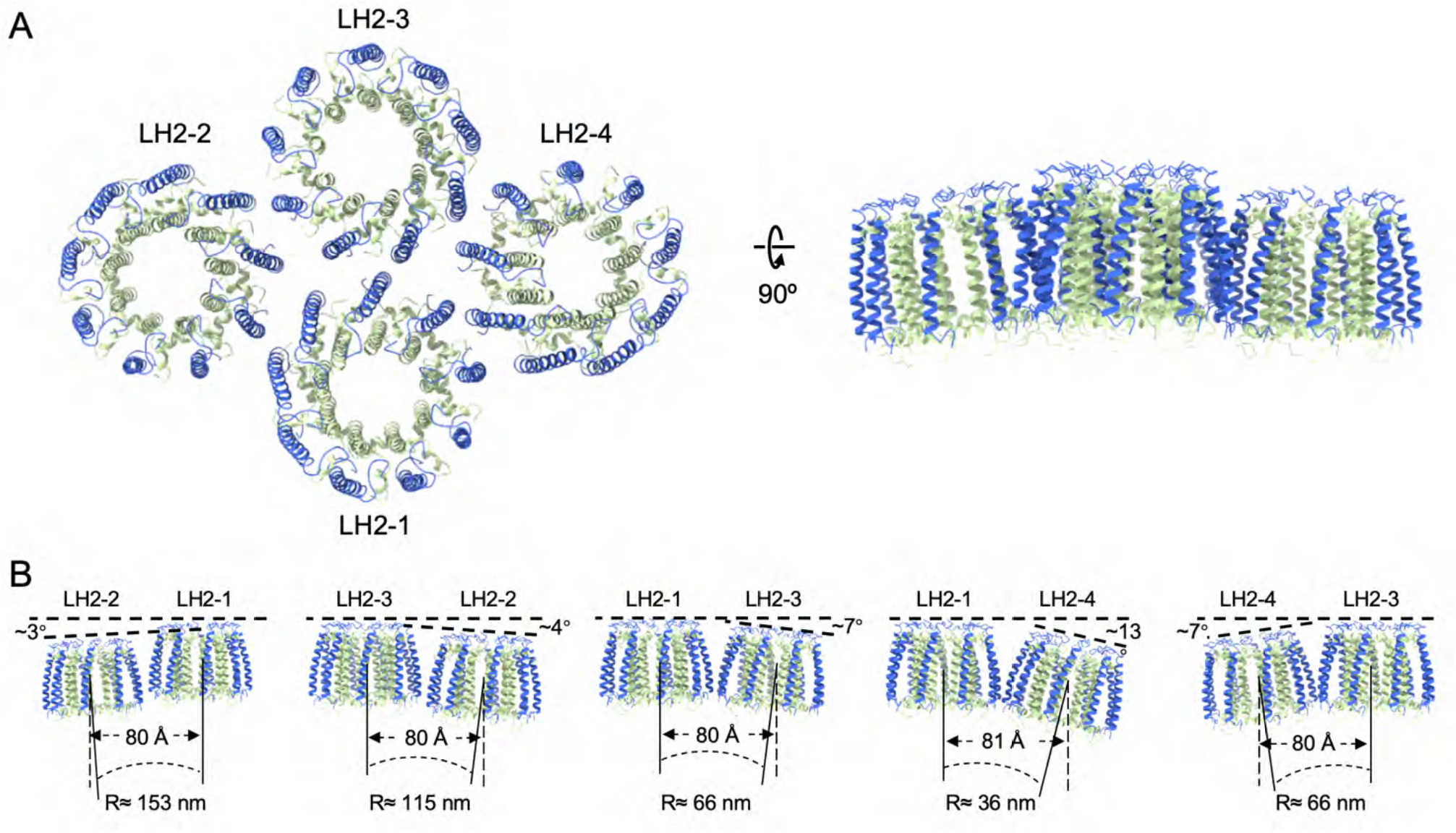
Structural organization of LH2-only assemblies. (A) Cytoplasmic and side views of the tetrameric LH2 assembly, with the four individual LH2 complexes labeled as LH2-1, LH2-2, LH2-3, and LH2-4. (B) Side views of pairwise LH2 assemblies, showing the relative angle between the two complexes, center-to-center distance, and curvature radius. The curvature radius was calculated according to the formula R = d/[2sin(θ/2)], where d is the center-to-center distance between adjacent complexes and θ is the relative angle between the two complexes.

AFM studies have visualized the PSU architecture and association between RC–LH1 and LH2 in photosynthetic membranes (10, 25), but lacked the resolution required to define their interactions. To our knowledge, the RC–LH1–LH2 and RC–LH1–2LH2 architectures represent the first cryo-EM structures showing how peripheral LH2 associates with the RC–LH1 core (Fig. 2A). In the RC–LH1– LH2 assembly, the primary interaction between RC–LH1 and LH2 is mediated by the LH1-2β subunit near the opening and the β subunit of LH2. The shortest distance between LH2 and LH1 subunits is 10.8 Å, and the closest center-to-center distance between BChls in the two complexes is 19.6 Å (Fig. 2A). Within the PSU, the RC–LH1 and LH2 complexes retain their individual architectures and associate through membrane-embedded interfaces with a tilt angle of 1.5° (Fig. 2B). This arrangement may be due to hydrophobic mismatch that can influence membrane-protein orientation, oligomerization, lateral organization, membrane-mediated protein–protein interactions, and membrane curvature (38, 39). Hydrophobic mismatch could also result in distinct surface protrusions of RC–LH1 and LH2, consistent with AFM observations of RC–LH1 and LH2 within photosynthetic membranes (40).

The larger RC–LH1–2LH2 assembly contains a second LH2 complex (LH2-2) positioned adjacent to the first LH2 complex (LH2-1), without remarkable structural rearrangements of the RC– LH1 core and LH2-1 (Fig. 2C). In this PSU architecture, the RC–LH1–LH2-1 interface resembles that observed in the RC–LH1–LH2 assembly. The newly formed interface between RC–LH1 and LH2-2 is mainly established through interactions between the LH1-4β subunit and the β subunit of LH2-2 (Fig. 2C). The two LH2 rings are separated by a center-to-center distance of 88 Å (Fig. 2C), and differ by ∼ 5° in rotational orientation (Fig. 2D). This nonrandom packing geometry illustrates how additional LH2 complexes can be incorporated into the RC–LH1–LH2 assembly without large organizational rearrangements, generating PSU assembly structures with variable antenna compositions.

### Curved LH2 oligomers extend the antenna network

Structural analysis of the tetrameric LH2 assembly reveals that adjacent LH2 complexes form a locally nonplanar cluster with relative tilts between neighboring complexes (Fig. 3A). Their relative intercomplex rotations of 3-13° correspond to local curvature radii of 36-153 nm, and the center-to-center distance of 80-81 Å between adjacent LH2 complexes defines a tightly packed antenna network (Fig. 3B). The experimentally observed rotations and corresponding curvature radii are in good agreement with the average tilt of 8.5 ± 0.3° and curvature radius of 49.5 nm predicted previously by computational simulations (41). Relative tilts of approximately 5° and 15° were also observed in reconstituted LH2 2D crystal membranes (42) and between parallel-oriented LH2–LH3 pairs reconstituted into nanodiscs (28), whereas the present tetramer reveals nonplanar antenna packing in an assembly isolated directly from photosynthetic membranes. The broader experimental range may capture flexibility in LH2 packing associated with membrane curvature, surface charge, or local packing constraints, whereas the earlier simulation likely represents an energetically favored average configuration (41).

### Excitation energy transfer within the photosynthetic unit

The isolated RC–LH1 and LH2 structures define pigment arrangements within each complex (Fig. 1), while the higher-order assemblies reveal how neighboring complexes associate (Figs. 2 and 3). Together, they provide a structural framework for modeling excitation migration within and between complexes.

In the RC–LH1–LH2 and RC–LH1–2LH2 assemblies, the LH2 complexes are positioned in direct contact with the LH1 ring, resulting in a minimum BChl-to-BChl separation of ∼20 Å between neighboring LH complexes (Figs. 2A, 2C). The close pigment proximity provides a structural basis for intercomplex EET. The neighboring complexes exhibit modest relative tilts of approximately 1.5° and 5° at the respective interfaces (Figs. 2B, 2D), maintaining an approximately coplanar arrangement of their pigment arrays. These features establish a continuous, unique BChl network connecting peripheral LH2 antennae to the RC–LH1 core.

To evaluate these pathways, we performed EET simulations using structural models derived from molecular dynamics simulations initiated from the experimental structures (Figs. 4, S11 and S12; Table S2). The EET dynamics were analyzed at two complementary levels. At the pigment level, Förster theory was used to characterize BChl-to-BChl energy transfer both within and between neighboring antenna complexes, including the B800→B850 and B800→B800 pathways and the fastest pigment-pair transfer pathways across intercomplex interfaces (Fig. 4 and Table S2). At the complex level, generalized Förster theory was employed to evaluate EET between the collective pigment manifolds of neighboring complexes (Fig. S12).

**Fig. 4.**
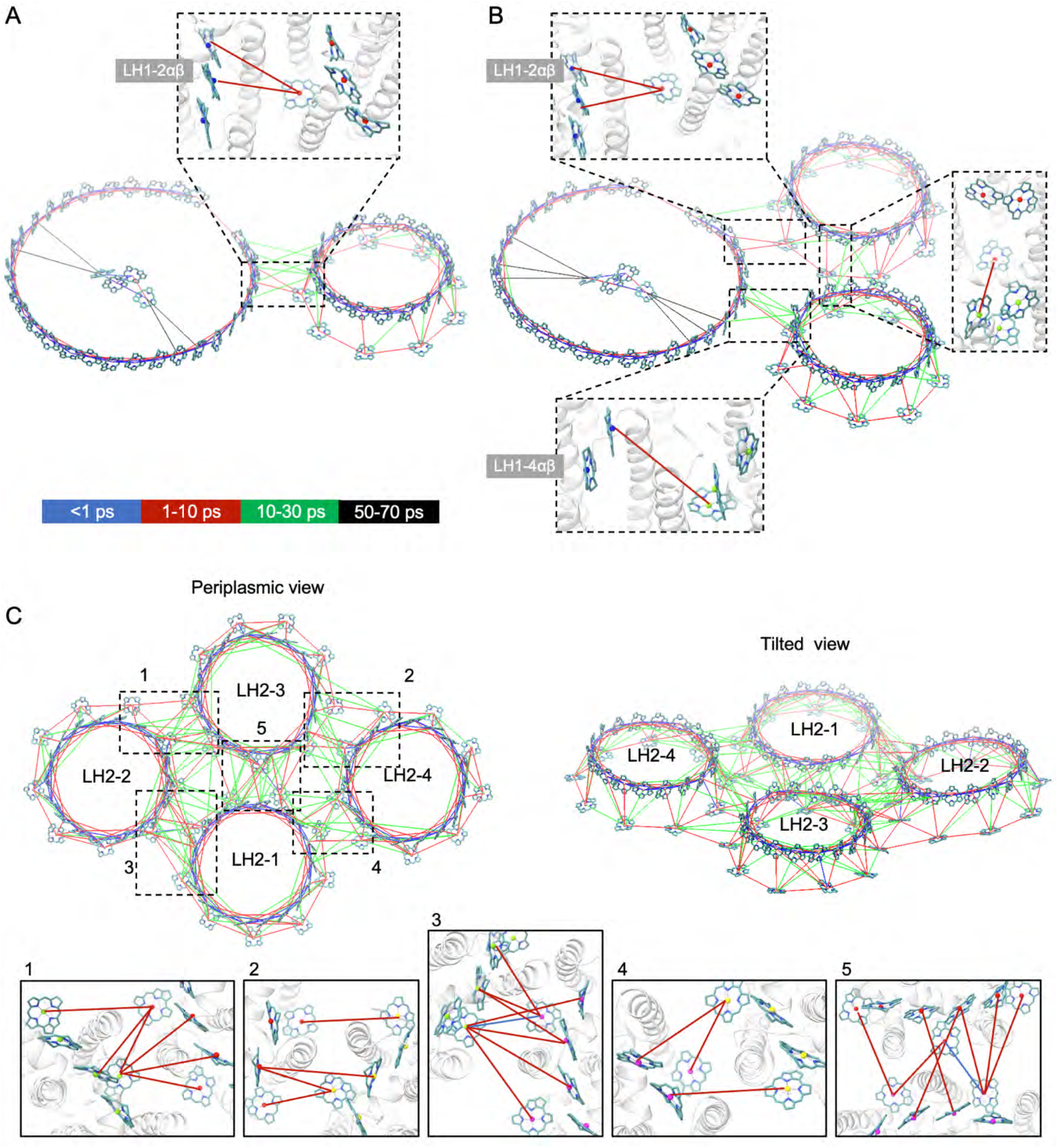
Excitation energy transfer pathways in supramolecular photosynthetic assemblies. (A) Energy-transfer network of the RC–LH1–LH2 assembly shown in tilted views. Intracomplex energy transfer dominates within the LH1 ring, whereas additional transfer pathways connect LH2 to the RC–LH1 core. The fastest pigment–pigment energy-transfer pathways between adjacent modules are indicated in the zoomed-in views. (B) Energy-transfer network of the RC–LH1–2LH2 assembly shown in tilted views, revealing continuous excitation-transfer pathways through adjacent LH2 complexes toward the RC–LH1 core. The fastest pigment–pigment energy-transfer pathways between adjacent modules are indicated in the zoomed-in views. (C) Energy-transfer network of the tetrameric LH2 assembly shown in cytoplasmic and tilted views. The fastest pigment–pigment energy-transfer pathways between adjacent modules are indicated in the zoomed-in views. Colored edges represent excitation-transfer events occurring on different timescales.

Within LH2, the calculated B800→B850 pathways yielded a mean transfer time of 1.7 ± 0.7 ps, broadly comparable to experimental results (43-45). B800→B800 transfer was slower, averaging 2.1 ± 0.8 ps, compared to the approximately 0.4-ps excitation redistribution detected by anisotropy measurements (44). The mean pairwise B850–B850 and B880–B880 transfer time constants were 0.018 and 0.016 ps, respectively, shorter than the approximately 0.09–0.30-ps timescales reported by ultrafast spectroscopy (46, 47). Although these pairwise estimates are not directly comparable to the collective excitation dynamics measured experimentally, these data are consistent with rapid EET between neighboring pigments in both antenna rings.

At the whole-complex level, transfer between adjacent LH2 complexes depended strongly on their packing geometry. The closely packed LH2 tetramer provides interconnected pathway for EET between neighboring LH2 antenna complexes (Fig. 4C). The five interfaces in the LH2 tetramer produced transfer times of 2.5–12.0 ps (Fig. S12), comparable to the ∼ 10-ps LH2→LH2 hopping time measured in photosynthetic membranes (48). Transfer between the two LH2 complexes associated with RC–LH1 was slower and direction-dependent, occurring within 16.7 and 19.0 ps. These calculations suggest faster energy transfer across several LH2 interfaces in the tetramer than between the two LH2 complexes in RC–LH1–2LH2.

For comparison, recent studies measured LH3→LH2 transfer times of 5.7, 9.8, and 14.7 ps in reconstituted nanodiscs with nearest intercomplex pigment separations of approximately 25, 28.5, and 31.4 Å, respectively (28). Simulations based on an AFM-derived chromatophore model estimated LH2→LH2 transfer times of 3–20 ps, with variations associated with the relative separation and geometry of neighboring complexes (49). Our calculated transfer times span a comparable picosecond regime and likewise indicate sensitivity to intercomplex geometry, although the different antenna compositions and experimental conditions preclude a direct quantitative comparison.

Calculated LH2→LH1 transfer times were 9.3 ps in RC–LH1–LH2 and 11.2 and 20.6 ps for the two nonequivalent LH2 antennae in RC–LH1–2LH2 (Fig. S12), broadly comparable to the approximately 10-ps timescales reported for membrane-associated and reconstituted systems (44, 48). The 20.6-ps pathway indicates comparatively weak energetic connectivity at one LH2–LH1 interface. Thus, association with the same RC–LH1 core does not confer equivalent EET kinetics on neighboring LH2 complexes; instead, their relative positions and pigment geometries modulate transfer to the core antenna.

Following excitation of LH1, transfer to the RC occurred within 28.3 ps in RC–LH1–LH2 and 35.9 ps in RC–LH1–2LH2 (Fig. S12). These values are comparable to experimentally reported values of approximately 35–50 ps for purple-bacterial RC–LH1 complexes (50, 51).

Collectively, structure-based simulations reproduce the established EET sequence from B800 to B850, between peripheral LH2 antennae, and subsequently through LH1 to the RC. They further predict that intracomplex redistribution is rapid, whereas intercomplex transfer is more sensitive to supramolecular packing, producing kinetically distinct pathways among PSUs with different antenna compositions.

To illustrate the organization of a large PSU, we generated a schematic model by aligning the RC–LH1–LH2 structure with the LH2 tetramer, yielding one RC–LH1 core associated with four LH2 complexes (Fig. 5A). Mapping the calculated transfer times onto this model outlines a stepwise pathway in which excitation equilibrates rapidly within LH2, migrates between adjacent LH2 complexes (∼3–12 ps), transfers to LH1 (∼10–20 ps), and subsequently reaches the RC (∼36 ps) (Fig. 5B). The model suggests that variations in LH2 packing could maintain a geometrically connected pigment network for energy migration toward the RC–LH1 core.

**Fig. 5.**
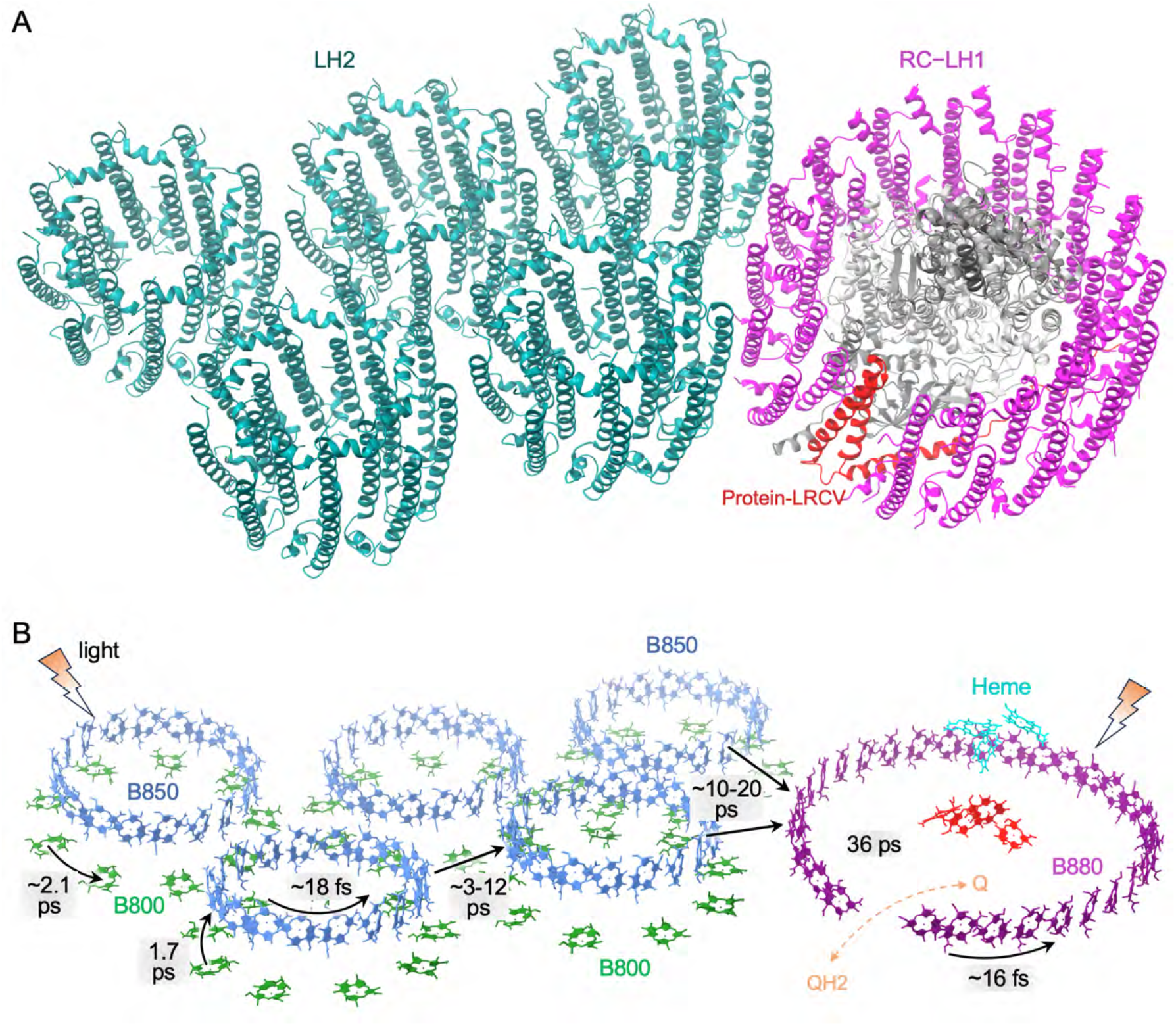
Proposed organization and excitation energy transfer pathways of a higher-order photosynthetic unit. (A) Tilted view of a composite model generated by superimposing the experimentally determined RC–LH1–LH2 and tetrameric LH2 assemblies through a shared LH2 complex, yielding one RC–LH1 core associated with four LH2 antennae. The composite illustrates a possible higher-order organization of the PSU rather than an experimentally resolved four-LH2 PSU. LH2, RC–LH1, and Protein-LRCV are indicated. (B) Proposed EET network mapped onto the composite model using calculations performed separately for the component assemblies. B800, B850, and B880 BChls are shown in green, blue, and purple, respectively; BChls of the RC are shown in red, and hemes are in cyan. Black arrows indicate selected excitation-transfer pathways, with representative calculated transfer time constants annotated. Intracomplex values describe selected pigment-pair transfers, whereas intercomplex values describe transfer between pigment domains. Orange dashed arrows schematically indicate quinone/quinol exchange (Q/QH²), potentially through the LH1 opening mediated by Protein-LRCV (which was not included in EET calculations). See also Fig. S12 and Table S2.

### Organizational modularity of bacterial photosynthetic units

The structural analysis and EET calculations provide insights into the organizational principle of bacterial PSUs. Despite differences in composition, the assembly maps accommodate the isolated RC–LH1 and LH2 structures without requiring large-scale rearrangements at the available resolution. The RC–LH1–LH2 and RC–LH1–2LH2 structures reveal different antenna–core organizational states (Fig. 2), while the LH2 tetramer extends this organizational framework to antenna–antenna networks (Fig. 3). Consistently, previous studies showed that LH2 can assemble independently and newly synthesized LH2 rings can be incorporated into core-rich membrane regions (22, 23, 26, 52-54). These observations support structural modularity of PSU assemblies.

The calculations further suggest that structurally distinct interfaces provide kinetically nonequivalent routes for excitation migration. EET between neighboring LH2 complexes is predicted to be faster within the tetramer than between the two core-associated LH2 antennae, while the nonequivalent LH2–LH1 interfaces yield different transfer times toward the same core (Figs. 4 and S12). Indeed, simulations indicate that intercomplex EET is dependent upon the packing of complexes in the photosynthetic membrane (49). Thus, variation in antenna organization can influence energetic connectivity even when the constituent complexes retain similar overall architectures.

Relative tilts and vertical offsets between neighboring complexes also produce nonplanar arrangements that may accommodate local membrane curvature (Figs. 2 and 3). These assemblies provide examples of how peripheral antennae can connect with one another and with the RC–LH1 core through different spatial arrangements, highlighting the organizational flexibility of photosynthetic modules. This packing could allow the constituent complexes to accommodate local membrane geometry and direct excitation migration and trapping in photosynthetic membranes.

Although these supercomplex assemblies were isolated directly from photosynthetic membranes, detergent solubilization and purification may selectively preserve stable associations while disrupting weaker contacts or altering intercomplex geometry, potentially leading to structural heterogeneity (Fig. S10). The recovered structures therefore provide examples of membrane-derived antenna–core and antenna–antenna organization, although the full diversity and relative abundance of these arrangements in native membranes remain to be investigated. Direct comparison with intact-membrane observations will help establish how closely the resolved interfaces reflect their native organization.

## Conclusion

To our knowledge, this study provides the first cryo-EM view of membrane-derived bacterial PSU assemblies comprising an intact RC–LH1 core associated with peripheral LH2 antennae. These higher-order assemblies reveal distinct antenna–core and antenna–antenna arrangements, in which RC–LH1 and LH2 complexes act as constituent modules, retaining their characteristic structures and associating through membrane-embedded interfaces with variable compositions. Differences in rotation, tilt, and vertical displacement suggest packing flexibility that may accommodate local membrane geometry while maintaining pigment connectivity between peripheral LH2 and the RC– LH1 core for efficient excitation transfer. Structure-based calculations predict picosecond intercomplex EET, with different transfer time constants at nonequivalent interfaces, suggesting that supramolecular geometry plays an important role in influencing energy migration toward the RC.

These findings provide insights into the organizational modularity and supramolecular packing of bacterial PSUs, which could mediate membrane architecture and tune light-harvesting capacity and energy transfer. More broadly, this work extends the structural understanding of the photosynthetic machinery from atomic structures of isolated pigment–protein complexes to their higher-order associations and provides a framework for investigating how membrane organization shapes photosynthetic energy flow and for guiding future efforts to engineer artificial photosynthesis.

## Materials and Methods

### Growth of *Rdv. viride*

Wild-type *Rhodovulum viride* (KCTC 15223) was obtained from the Korean Collection for Type Cultures (KCTC). *Rdv. viride* cells were grown phototrophically under anaerobic conditions in liquid 2216E medium at 30°C in glass bottles under a light intensity of 25 μmol photons m^™2^ s^™1^ (Bellight 70 W halogen bulbs).

### Purification of RC–LH1 and LH2 samples

Cells were harvested by centrifugation at 5,000 × *g* for 10 min at 4°C, washed three times with Tris-HCl (pH 8.0), and resuspended in 20 mM HEPES (pH 8.0). The cells were disrupted by passage through a French press three times at 16,000 psi. Cell debris was removed by centrifugation at 20,000 × *g* for 30 min. Membranes were collected by centrifuging the resulting supernatant at 125,000 × *g* for 90 min and solubilized by adding β-DDM (n-dodecyl β-D-maltoside) to a final concentration of 3% (w/v) for 30 min to 60 min in the dark at 4°C with gentle stirring. Unsolubilized proteins were removed by centrifugation at 21,000 × *g* for 30 min. The supernatant was then applied to 10–25% (w/v) continuous sucrose gradients prepared with working buffer containing 0.01% (w/v) β-DDM. Gradients were centrifuged at 230,000 × *g* for 18 h. The RC–LH1 and LH2 complexes in the sucrose gradient solution were collected, and the purity of RC–LH1 and LH2 complexes was characterized by sodium dodecyl sulfate-polyacrylamide gel electrophoresis (SDS-PAGE) and absorption spectra (Fig. S1).

### Absorption spectroscopy

Absorbance spectra of purified RC–LH1 and LH2 samples were recorded from 300 to 900 nm at 1-nm intervals at room temperature using a Libra S22 spectrophotometer (Biochrom, United Kingdom).

### Bioinformatics analysis

Protein-LRCV from *Rdv. viride* and protein-3h from *Rdv. sulfidophilum* were obtained from their annotated genome sequences available in the NCBI Genome database. The sequences were aligned using Clustal Omega (v1.2.4) and visualized using ESPript (v3.2) (https://espript.ibcp.fr/ESPript/ESPript/).

### Cryo-EM data collection

Aliquots of 4 μL RC–LH1 and LH2 samples were separately applied to freshly glow-discharged holey carbon grids (Quantifoil Au R2/1, 200 mesh) with a continuous carbon support. The grids were blotted for 2 s at 100% humidity and 10°C, with a blotting force level of 0, and immediately plunge-frozen into liquid ethane cooled by liquid nitrogen using a Vitrobot Mark IV (Thermo Fisher Scientific, USA). The grids were then loaded into a 200 kV Glacios 2 microscope (Thermo Fisher Scientific, USA) that was equipped with a Falcon 4 direct electron detector (Thermo Fisher Scientific, USA) for data acquisition. A total of 5,962 movie stacks were automatically recorded using EPU (Thermo Fisher Scientific, USA) (55) at a total dose of 40 e^-^/Å^2^ per stack, with a defocus range of -0.8 to -1.8 μm and a pixel size of 0.89 Å.

### Data processing

All cryo-EM data processing was performed using cryoSPARC (v4.4.1). Movie stacks were subjected to patch motion correction and contrast transfer function (CTF) estimation, and micrographs with an estimated CTF resolution worse than 7 Å were discarded (Table S1). For the RC–LH1 sample, particles were picked using the Blob picker algorithm and the Template picker algorithm in cryoSPARC. After particle extraction and 2D classification, high-quality particles were selected for further analysis. These particles were used for ab initio reconstruction and subsequent 3D classification. The selected 3D classes with the best density features were then subjected to non-uniform refinement in cryoSPARC. This procedure yielded density maps corresponding to the RC– LH1, RC–LH1–2LH2, RC–LH1–LH2, and LH2 tetramer complexes, with resolutions of 2.58 Å, 6.77 Å, 5.68 Å, and 7.72 Å, respectively (Fig. S2). For the LH2 sample, particles were picked using the Blob picker algorithm in cryoSPARC. After particle extraction and 2D classification, high-quality particles were selected for further analysis. These particles were used for ab initio reconstruction and subsequent 3D classification. The selected 3D classes with the best density features were then subjected to non-uniform refinement in cryoSPARC. The final LH2 density map was determined at a resolution of 2.87 Å (Fig. S3). All resolution estimates were based on the gold-standard Fourier shell correlation (FSC) criterion at 0.143.

### Model building and refinement

The RC–LH1 complex structure of *Rdv. sulfidophilum* (PDB ID: 9WQV) and the LH2 complex structure of *Rba. sphaeroides* (PDB ID: 7PBW) were initially docked into the cryo-EM density maps of the RC–LH1 and LH2 complexes, respectively, using UCSF Chimera (v1.17) (56). The models were manually revised and refined based on cryo-EM density using Coot (v0.9.4) (57), followed by real-space refinement using Phenix (v1.20.1) (58). To construct models of the higher-order assemblies, the high-resolution RC–LH1 and LH2 structures determined in this study were individually docked into the experimentally determined density maps of the RC–LH1–LH2, RC–LH1–2LH2, and LH2 tetrameric complexes. Given the moderate-to-low resolutions of these maps, the final models were largely retained from the docked components without further detailed refinement (59, 60). Figures were generated using UCSF Chimera (v1.17) and ChimeraX (v1.16) (61).

### Molecular dynamics (MD) simulations

Three photosynthetic assemblies, including the LH2 tetramer, RC–LH1–LH2, and RC–LH1–2LH2, were constructed based on the cryo-EM structures. The complexes were positioned in POPC lipid bilayers using CHARMM-GUI (62, 63) and VMD (64), solvated with TIP3P water (65), and neutralized with NaCl to a concentration of 0.15 M. All-atom MD simulations were performed using NAMD 2.13 (66) with the AMBER ff14SB force field for proteins (67) and Lipid17 for POPC. Parameters for bacteriochlorophyll (BChl) and bacteriopheophytin (BPhe) were adapted from previously reported parameters for bacterial photosynthetic cofactors (68), whereas neurosporene (SPO) and ubiquinone-10 (U10) were parameterized using the GAFF force field (69). RESP charges were used where applicable (70), and missing hydrogen atoms were added using TLEAP (71). Long-range electrostatic interactions were treated using the particle-mesh Ewald method (72) with a grid spacing of approximately 1 Å, and a cutoff of 10 Å was applied to nonbonded interactions. Bonds involving hydrogen atoms were constrained using SHAKE (73), allowing an integration time step of 2 fs. Following energy minimization and gradual heating to 300 K, each system was equilibrated for 1 ns under NVT conditions, followed by 10 ns of restrained NPT equilibration. The restraints were then removed, and each of the three systems was subjected to a 500 ns NPT production simulation at 300 K and 1 atm using Langevin dynamics for temperature control and the Langevin piston method (74) for pressure control.

### Quantum Mechanics (QM) / Molecular Mechanics (MM) calculations and excitonic parameters

The final snapshot of each 500 ns MD trajectory was used for subsequent QM/MM calculations. Geometry optimization of individual BChl pigments and their local environments was performed using a modified version of pDynamo (75) at the AM1/MM level with electrostatic embedding (76, 77). The QM region comprised the BChl chromophore and protein residues within 3 Å of the BChl that coordinate or directly interact with the pigment. The phytol chain was truncated after the ester group, and the QM/MM covalent boundary was treated using link atoms. Molecules within 7 Å of the QM region were included in the movable MM region, whereas the remaining environment within 40 Å of the pigment was retained with fixed coordinates. Based on the optimized structures, vertical excitation energies of individual BChls were calculated at the TD-CAM-B3LYP/6-31G* level (78) within the electrostatic embedding of the surrounding MM environment using Gaussian 16 (79). Electronic couplings between pigments were evaluated using the transition-charge from electrostatic potential (TrEsp) approach (80, 81), with transition charges generated using Multiwfn (82).

### Excitation energy transfer calculations

The pigment network was described using a Frenkel exciton Hamiltonian constructed from the calculated site energies and electronic couplings (83). EET rates between individual pigments were calculated using Förster resonance energy-transfer theory (84, 85), with spectral overlap determined from normalized donor emission and acceptor absorption spectra represented by Gaussian line shapes (86). For energy transfer between pigment domains, generalized Förster theory was employed as reported previously (87-90), with donor and acceptor domains defined according to the naturally occurring pigment–protein structural units resolved in the cryo-EM structures. The calculated transfer rate *k*^*mn*^ was converted to the corresponding characteristic transfer time according to *τ*^*mn*^ = 1/*k*^*mn*^, and these transfer times were used to construct the EET networks and characterize the major excitation-transfer pathways among LH2, LH1, and RC pigments. The spectral parameters and environmental screening treatment followed our previously established protocol (89, 90).

