## Supplemental figures and tables for "Architecture and Energy Transfer of the Bacterial Photosynthetic Unit"

**Supplementary Materials**

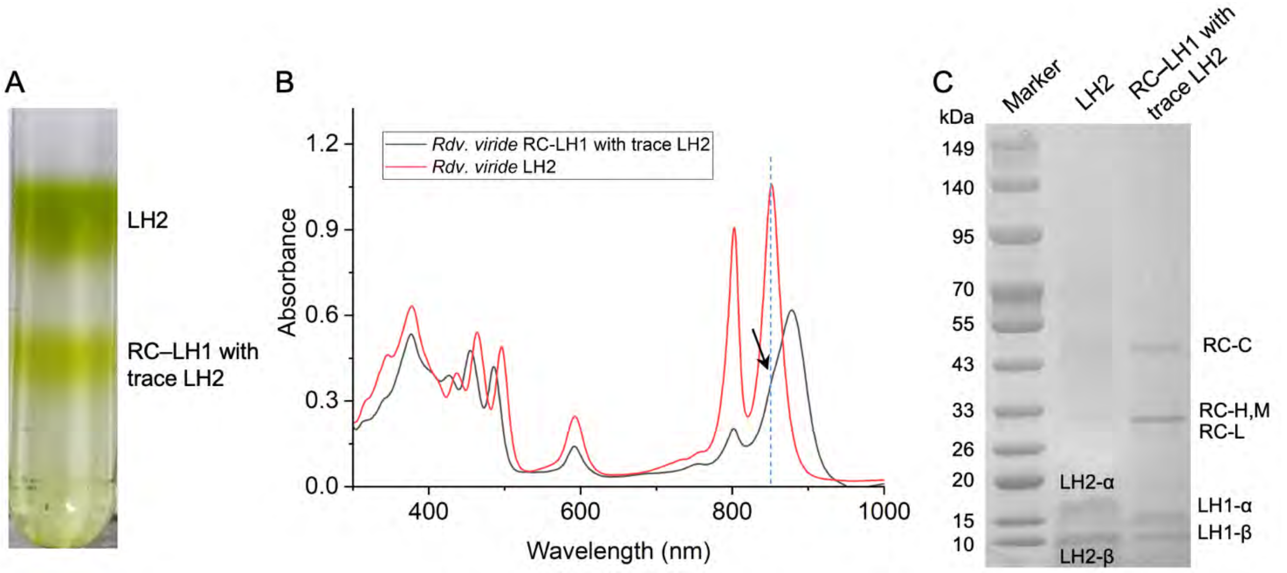


**Fig. S1. Purification and characterization of *Rdv. viride* RC–LH1 and LH2.** (A) Sucrose gradient ultracentrifugation of photosynthetic membrane complexes from *Rdv. viride*. (B) Room-temperature UV–vis absorption spectra of isolated LH2 and RC–LH1-enriched fractions. The arrowed shoulder peak of RC–LH1 with trace LH2 overlaps with the LH2 absorption peak at 850 nm. (C) SDS-PAGE analysis of the purified samples.

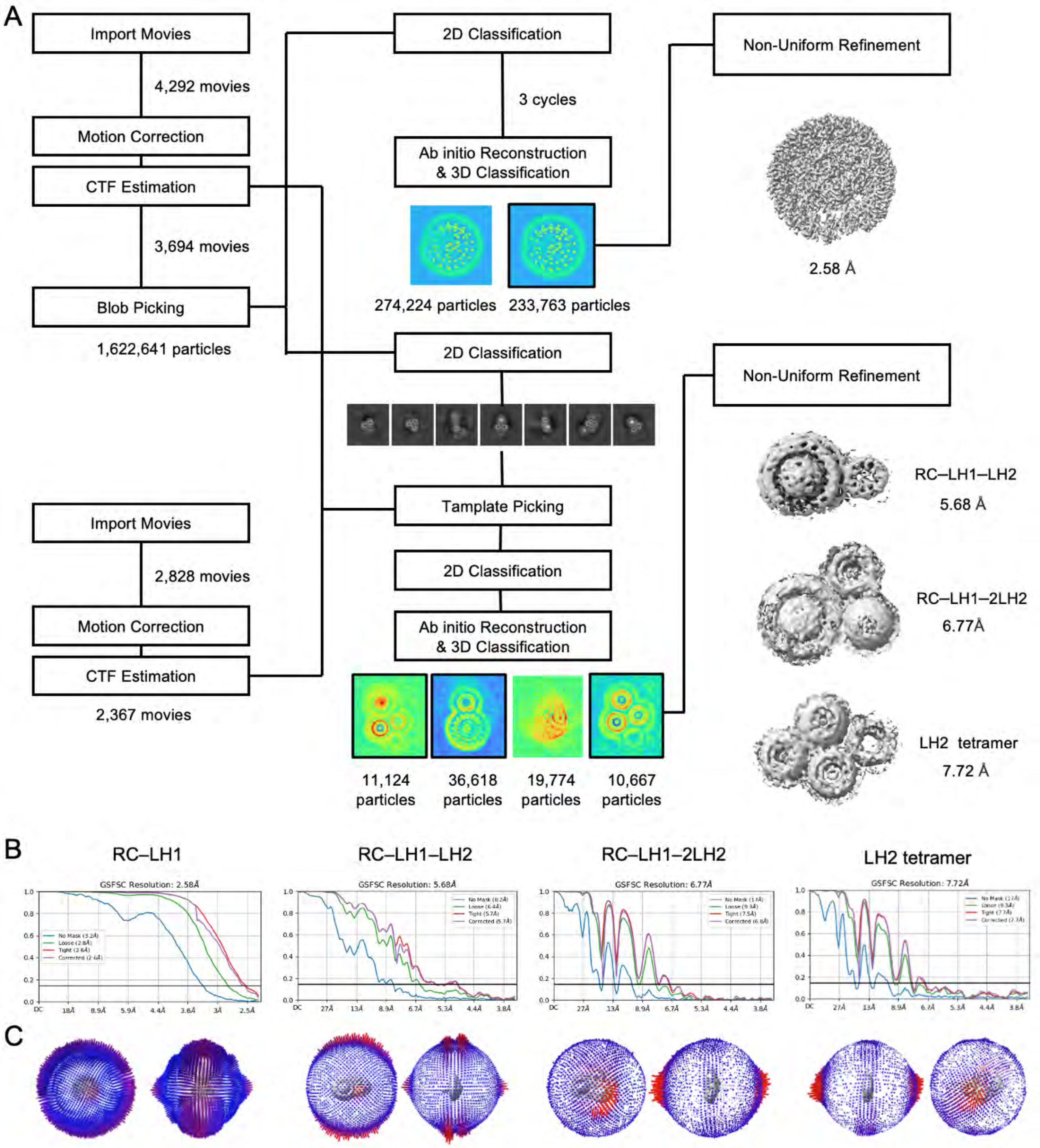


**Fig. S2. Cryo-EM data processing of *Rdv. viride* RC–LH1 containing trace LH2.** (A) Overview of cryo-EM data processing. (B) Fourier shell correlation (FSC) curves generated by cryoSPARC. Global resolution values were calculated according to the gold-standard FSC = 0.143. (C) Angular distributions of the anti-parallel and parallel orientations of particles contributing to the RC–LH1, RC–LH1–LH2, RC–LH1–2LH2, and LH2 tetramer reconstructions.

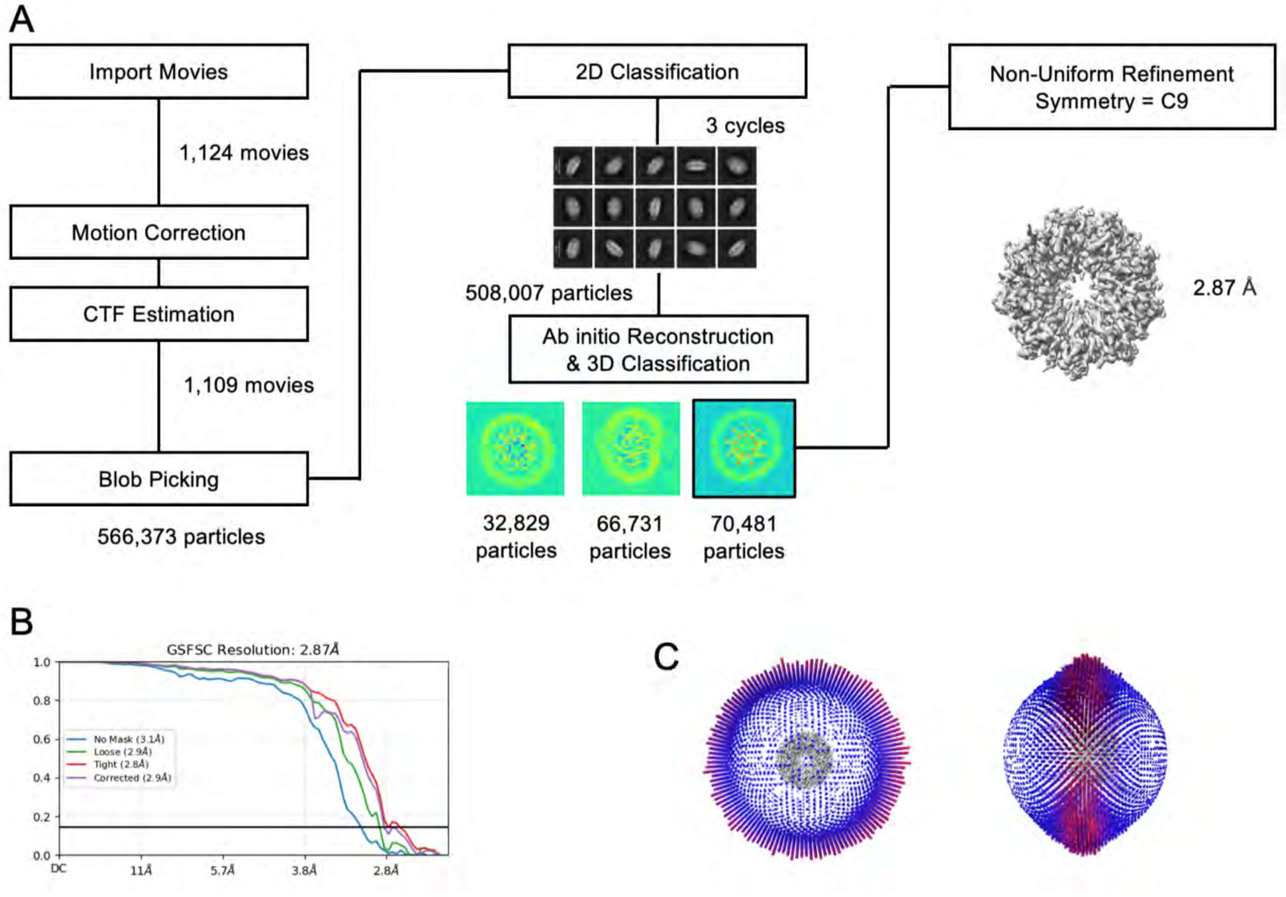


**Fig. S3. Cryo-EM data processing of *Rdv. viride* LH2.** (A) Overview of cryo-EM data processing. (B) Fourier shell correlation (FSC) curves generated by cryoSPARC. Global resolution values were calculated using the gold-standard FSC threshold of 0.143. (C) Angular distributions of the anti-parallel and parallel orientations of particles contributing to the LH2 reconstruction.

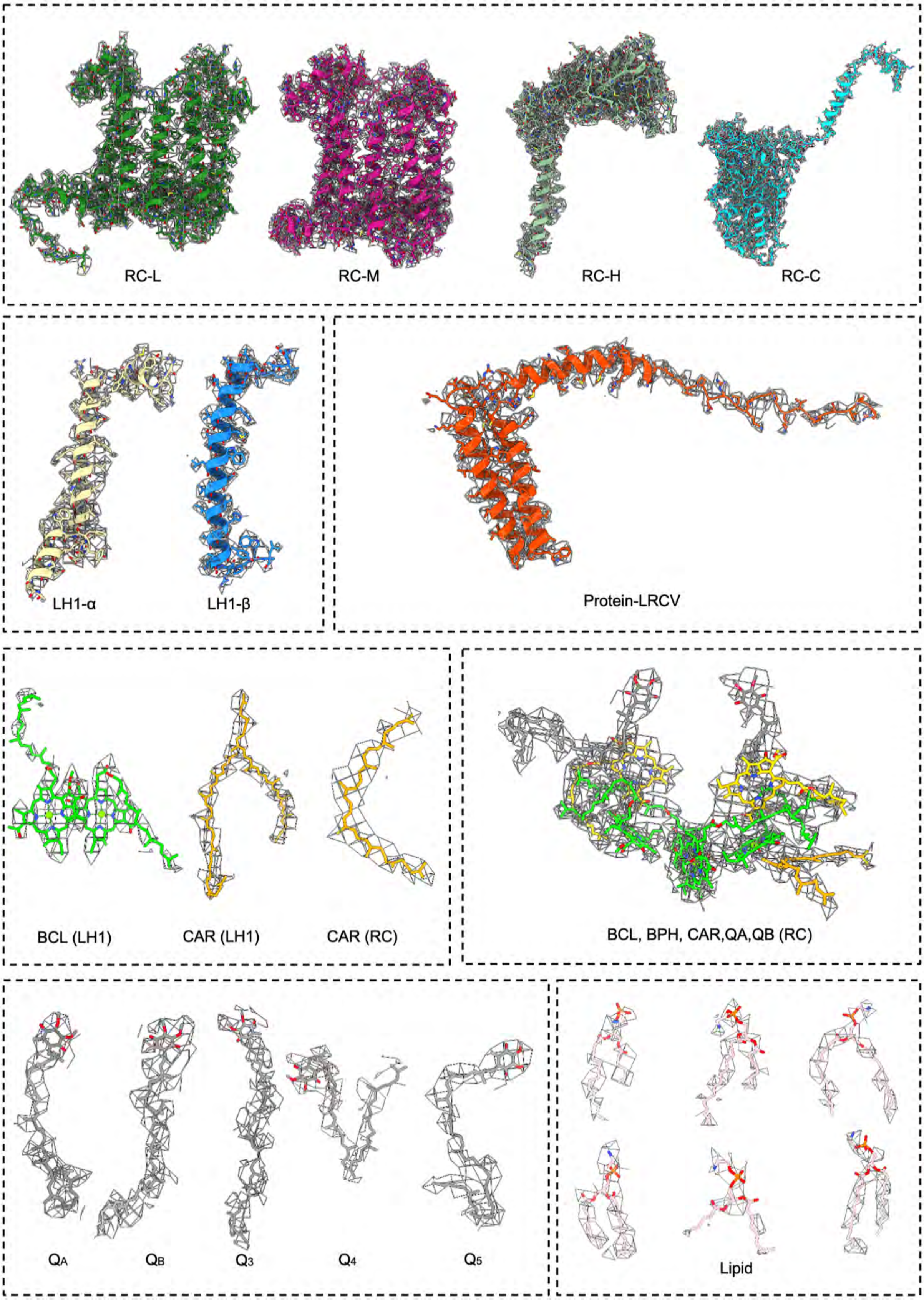


**Fig. S4. Cryo-EM map densities and structural models of protein regions and cofactors in the *Rdv. viride* RC–LH1 complex.**

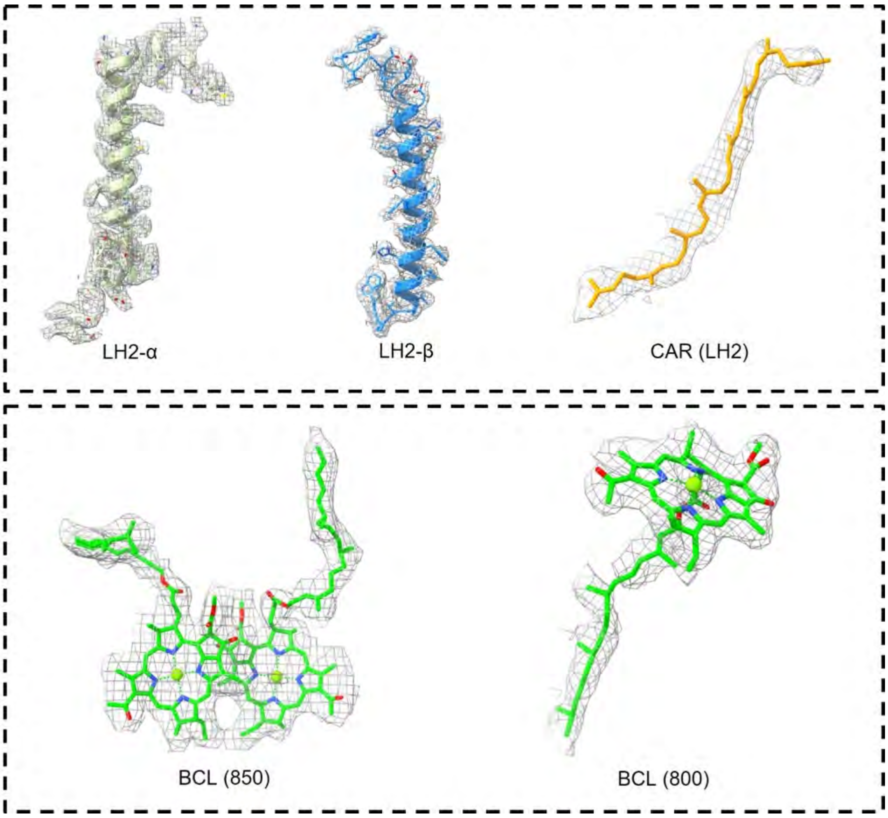


**Fig. S5. Cryo-EM map densities and structural models of proteins and cofactors in the *Rdv. viride* LH2 complex.**

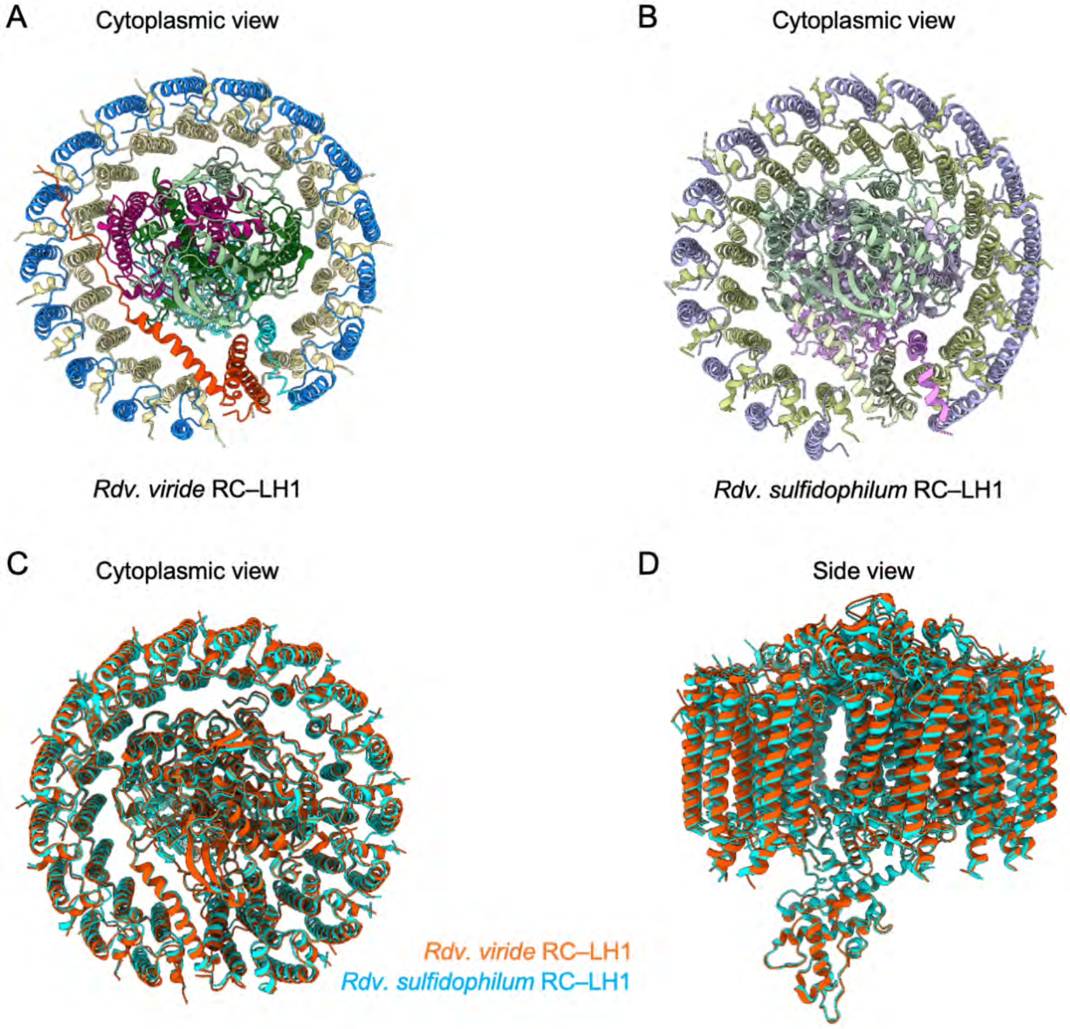


**Fig. S6. Structural comparison of the RC–LH1 complexes from *Rdv. viride* and *Rdv. sulfidophilum* (9WQV).** (A) Cytoplasmic view of the *Rdv. viride* RC–LH1 complex. (B) Cytoplasmic view of the *Rdv. sulfidophilum* RC–LH1 complex. (C, D) Structural comparison of the *Rdv. viride* and *Rdv. sulfidophilum* RC–LH1 complexes.

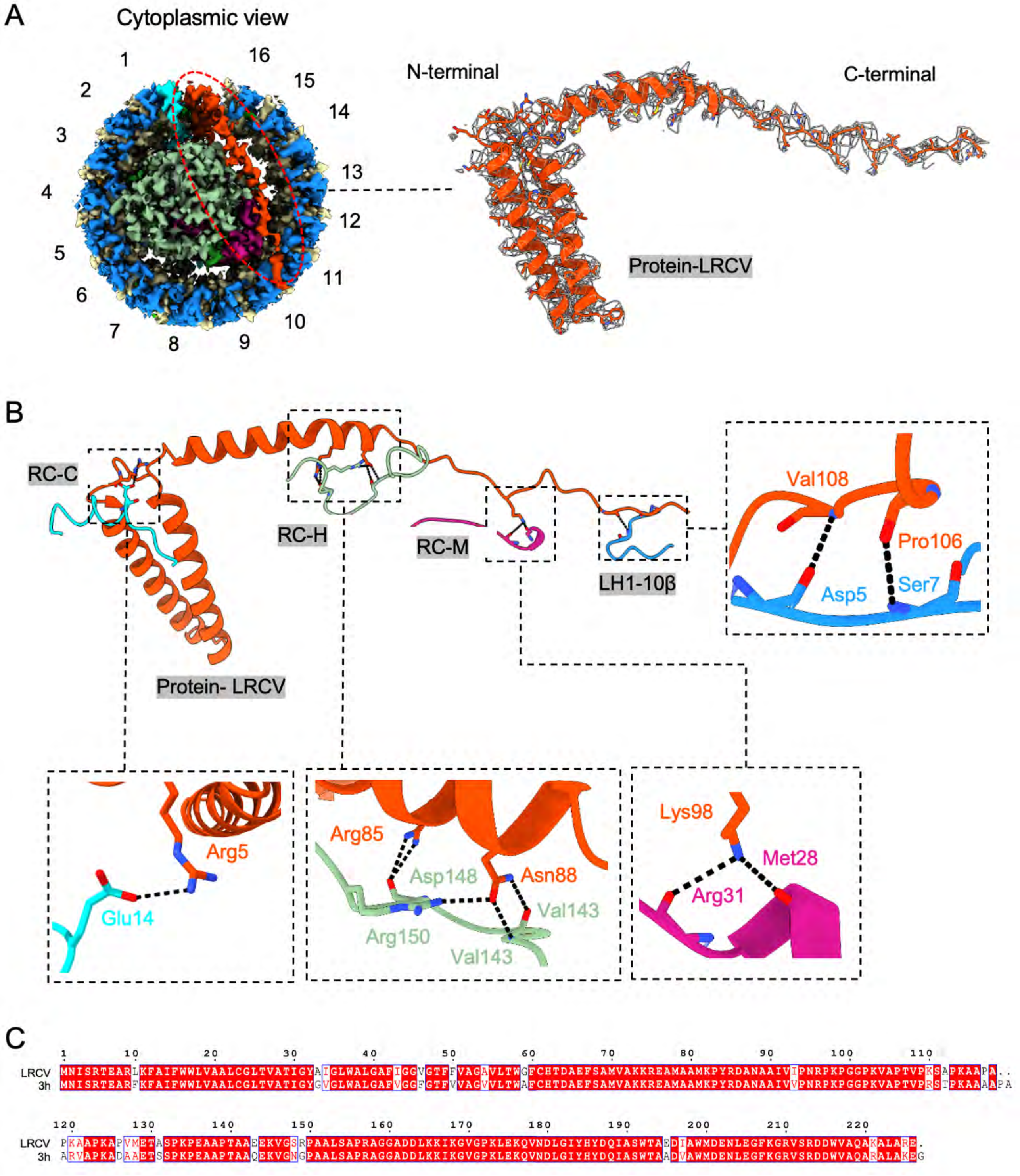


**Fig. S7. Structural analysis of protein subunit Protein-LRCV.** (A) Binding site of the Protein-LRCV subunit and the corresponding cryo-EM density map. (B) Interactions between the Protein-LRCV subunit and the surrounding subunits. Detailed interactions are shown in zoomed-in views. (C) Sequence alignment of Protein-LRCV from *Rdv. viride* and protein-3h from *Rdv. sulfidophilum* (PDB ID: 9WQV).

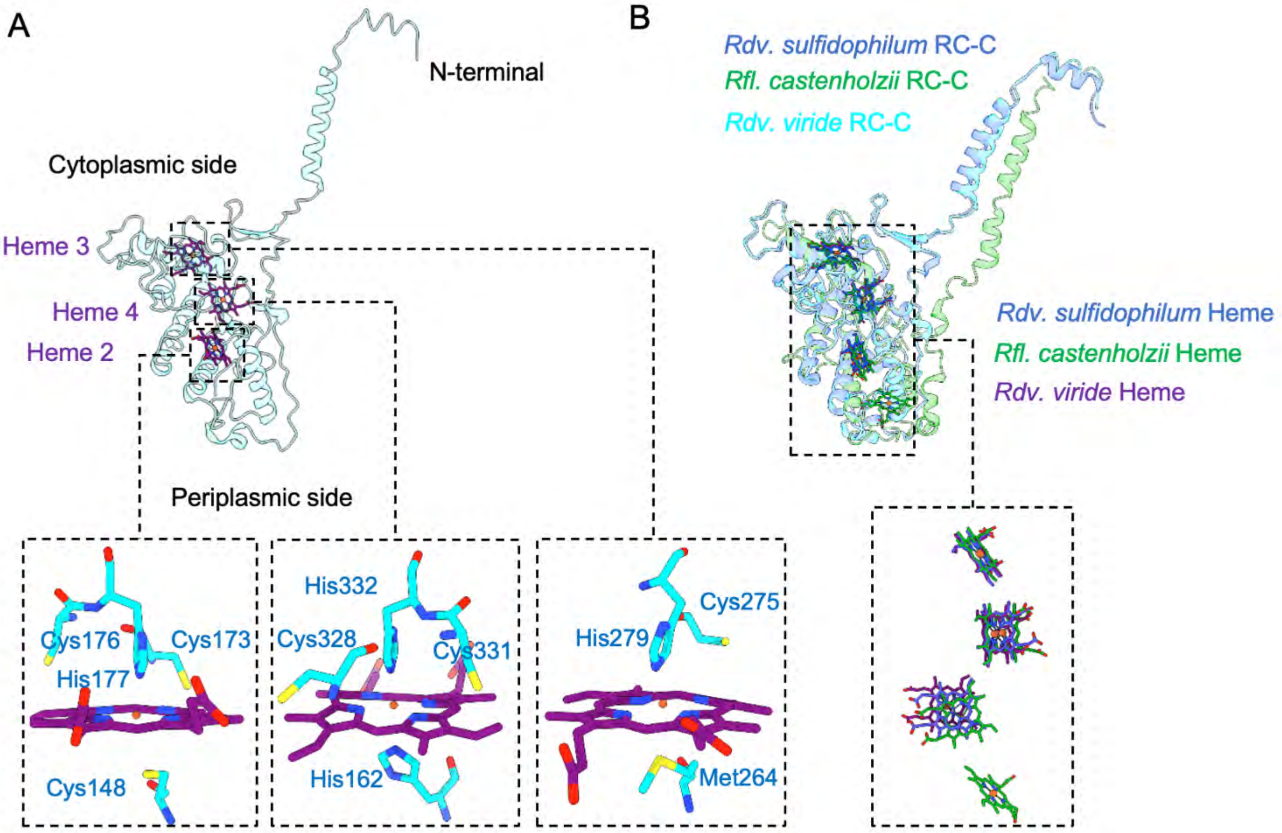


**Fig. S8. Structural analysis of the cytochrome c subunit.** (A) Arrangement of the heme groups in the cytochrome c subunit and their interactions with adjacent residues. Detailed interacting environments surrounding the hemes are shown in zoomed-in views. (B) Structural comparison of the cytochrome c subunits from *Rdv. viride*, *Rdv. sulfidophilum* (9WQV), and *Roseiflexus* (*Rfl.*) *castenholzii* (8IUG).

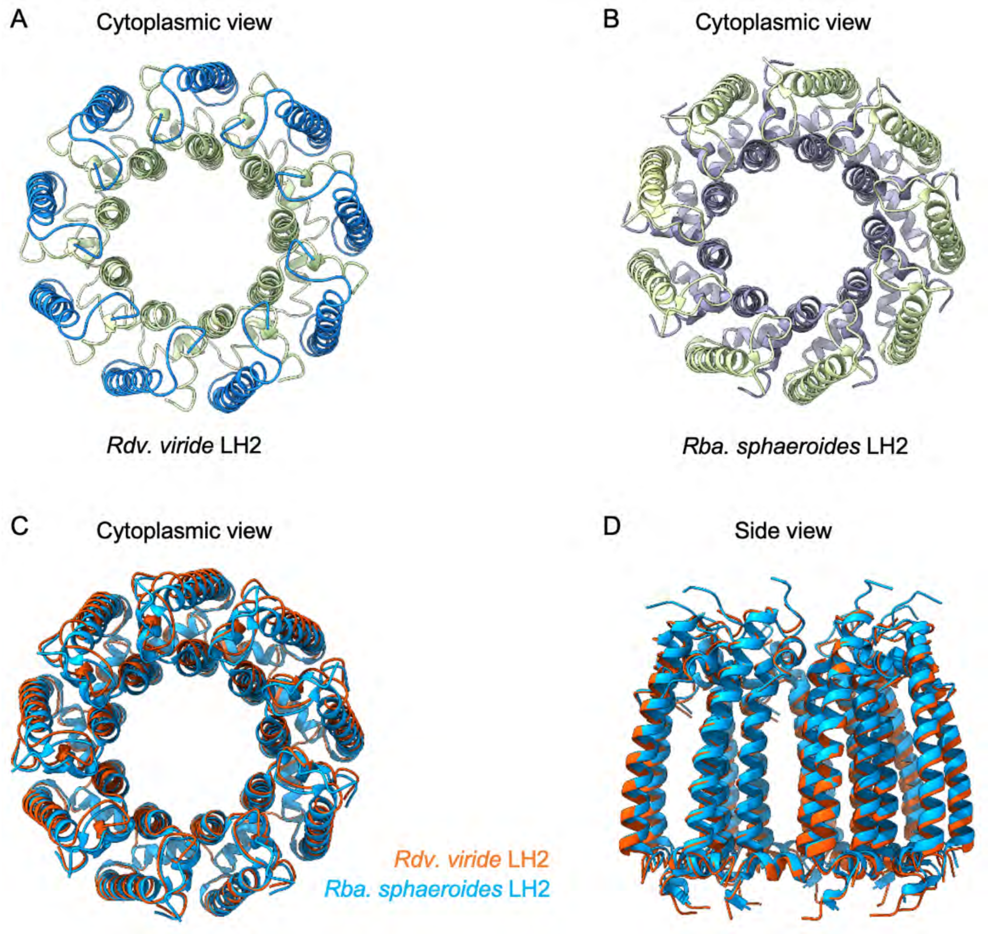


**Fig. S9. Structural comparison of the LH2 complexes from *Rdv. viride* and *Rba. sphaeroides* (7PBW).** (A) Cytoplasmic view of the *Rdv. viride* LH2 complex. (B) Cytoplasmic view of the *Rba. sphaeroides* LH2 complex. (C, D) Structural comparison of the *Rdv. viride* and *Rba. sphaeroides* LH2 complexes.

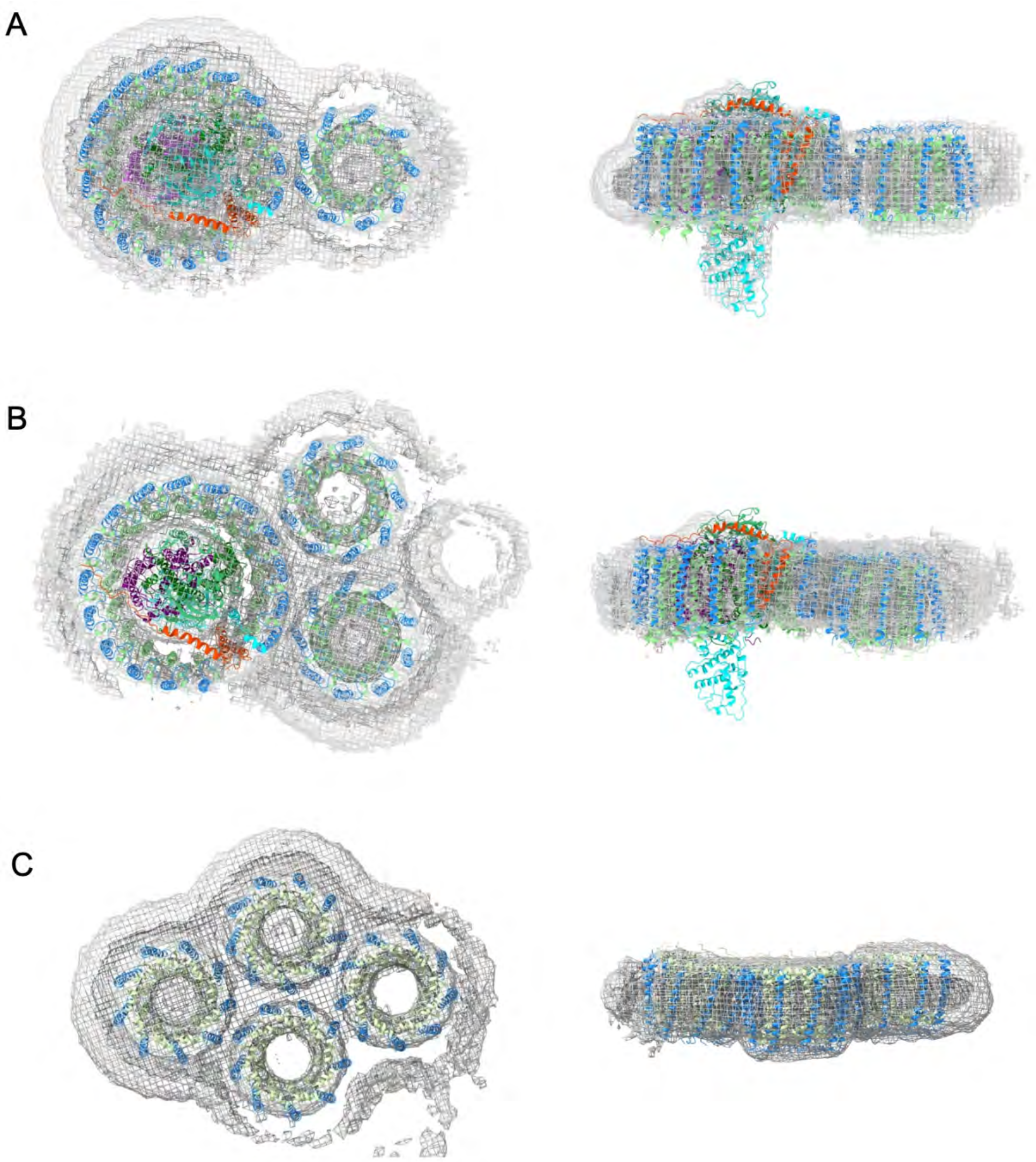


**Fig. S10. Cryo-EM map density and structural model overlays of RC–LH1–LH2 (A), RC–LH1–2LH2 (B), and the LH2 tetramer (C).**

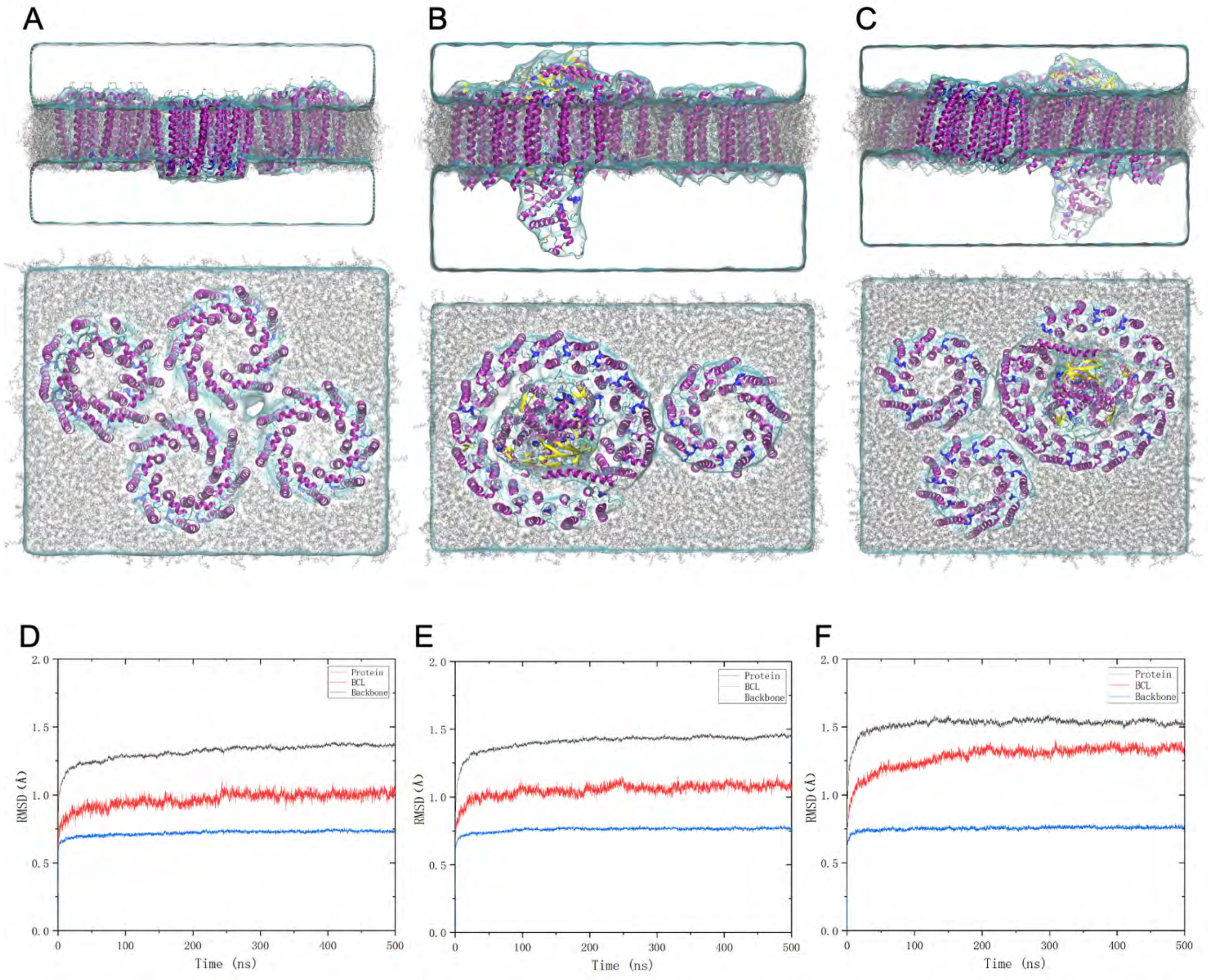


**Fig. S11. Molecular dynamics simulation systems and structural stability of the purple bacterial photosynthetic complexes.** (A–C) Representative structures of the LH2 tetramer, RC–LH1–LH2, and RC–LH1–2LH2 systems, respectively, embedded in a POPC lipid bilayer and solvated with water. For each system, side (top) and top (bottom) views are shown to illustrate the membrane embedding and spatial organization of the complexes. Protein secondary structures are shown in cartoon representation, POPC molecules in lines, and cofactors and pigments in licorice representation. (D–F) Time evolution of the root-mean-square deviation (RMSD) during the 500 ns MD simulations of the corresponding LH2 tetramer, RC–LH1–LH2, and RC–LH1–2LH2 systems. RMSDs are shown for all protein atoms (black), BChl molecules (BCL, red), and the protein backbone (blue). The RMSD profiles reach stable plateaus after the initial structural relaxation, indicating that the overall protein architectures and embedded BChl pigments remain conformationally stable over the subsequent simulation period.

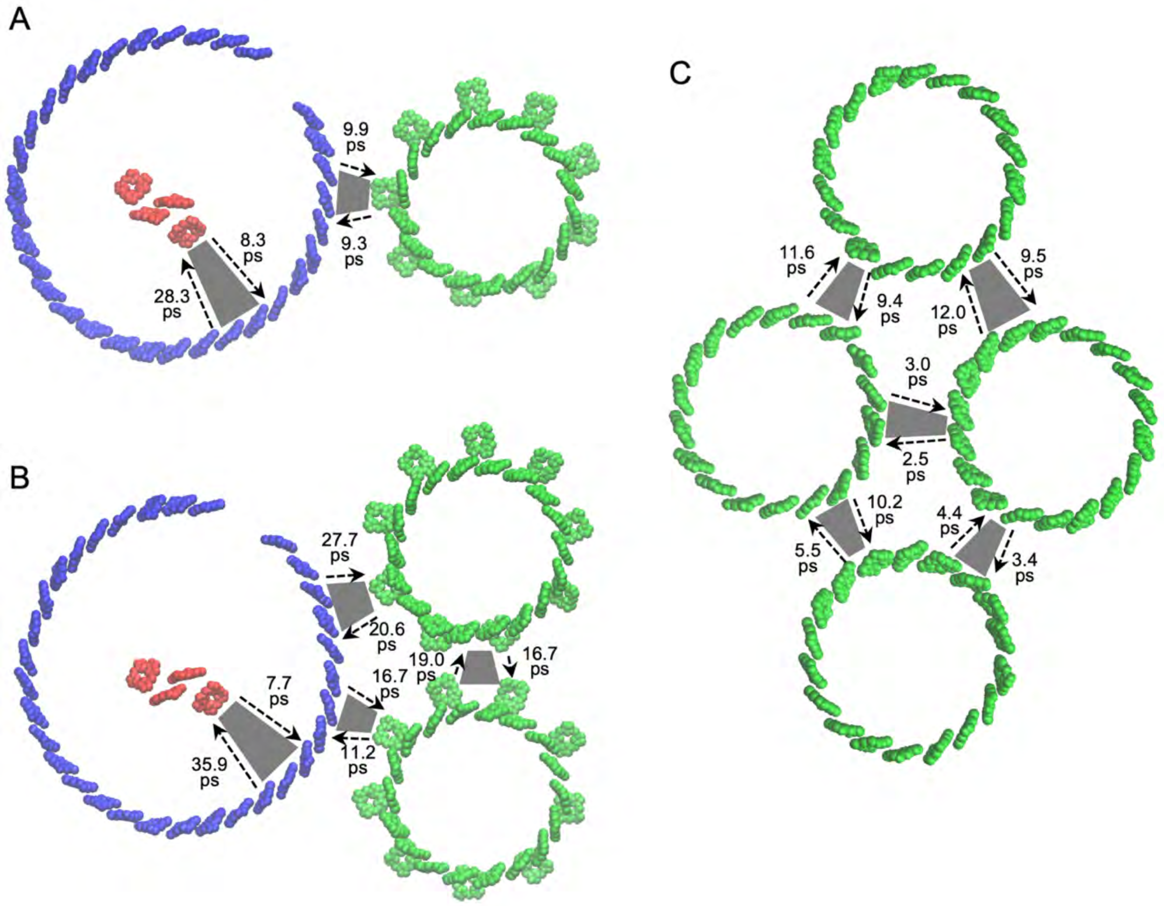


**Fig. S12. Generalized Förster theory (gFRET)-calculated EET timescales between neighboring photosynthetic complexes in the RC–LH1–LH2 assembly (A), RC–LH1–2LH2 assembly (B), and tetrameric LH2 assembly (C).** Arrows indicate the directions of EET, and the corresponding transfer timescales are given in ps.

**Table S1. Cryo-EM data collection, reconstruction, and model refinement statistics.**

| | RC–LH1 | LH2 | RC–LH1–LH2 | RC–LH1–2LH2 | LH2 tetramer |
|---|---|---|---|---|---|
| **Data collection and processing** | | | | | |
| Magnification (nominal) | 130,000 | 130,000 | 130,000 | 130,000 | 130,000 |
| Voltage (kV) | 200 | 200 | 200 | 200 | 200 |
| Electron exposure ($e^-/Å^2$) | 40 | 40 | 40 | 40 | 40 |
| Camera | FEI Falcon 4 | FEI Falcon 4 | FEI Falcon 4 | FEI Falcon 4 | FEI Falcon 4 |
| Defocus range (μm) | -0.8 ~ -1.8 | -0.8 ~ -1.8 | -0.8 ~ -1.8 | -0.8 ~ -1.8 | -0.8 ~ -1.8 |
| Pixel size (Å) | 0.89 | 0.89 | 0.89 | 0.89 | 0.89 |
| Symmetry imposed | C1 | C9 | C1 | C1 | C1 |
| Final particle images (no.) | 233,763 | 70,481 | 36,618 | 11,124 | 10,667 |
| FSC threshold | 0.143 | 0.143 | 0.143 | 0.143 | 0.143 |
| Map resolution (Å) | 2.58 | 2.87 | 5.68 | 6.77 | 7.72 |
| **Refinement** | | | | | |
| Initial model used (PDB code) | 9WQV | 7PBW | *Rdv. viride* RC–LH1 and LH2 | *Rdv. viride* RC–LH1 and LH2 | *Rdv. viride* LH2 |
| Model composition | | | | | |
| Non-hydrogen atoms | 27241 | 8631 | 35872 | 44503 | 34524 |
| Protein residues | 2856 | 846 | 3702 | 4548 | 3384 |
| Bfactor ($Å^2$) | | | | | |
| Protein | 44.09 | 20.57 | 45.09 | 45.68 | 48.32 |
| Ligand | 39.41 | 16.70 | 39.13 | 38.49 | 36.52 |
| R.m.s deviations | | | | | |
| Bond lengths (Å) | 0.013 | 0.006 | 0.013 | 0.014 | 0.015 |
| Bond angles (°) | 1.310 | 0.964 | 1.450 | 1.727 | 2.075 |
| Validation | | | | | |
| MolProbity score | 1.55 | 1.33 | 1.53 | 1.52 | 1.61 |
| Poor rotamers (%) | 0.47 | 0 | 0.37 | 0.30 | 0.19 |
| Ramanchandran plot | | | | | |
| Favored (%) | 97.95 | 97.78 | 98.30 | 98.41 | 98.98 |
| Allowed (%) | 2.01 | 2.22 | 1.67 | 1.57 | 0.99 |
| Disallowed (%) | 0.04 | 0 | 0.013 | 0.02 | 0.03 |
| PDB accession code | 43YS | 44VH | 43YV | 43YW | 43YX |
| EMDB accession code | EMD-82397 | EMD-82988 | EMD-82399 | EMD-82400 | EMD-82401 |

**Table S2. EET time constants calculated between selected BChl pairs within photosynthetic complexes using Förster resonance energy transfer (FRET) theory.**

| Energy transfer pathway | Minimum (ps) | Maximum (ps) | Mean ± SD (ps) | *n* |
|---|---|---|---|---|
| LH2 B800-B800 | 1.1 | 4.8 | 2.1 ± 0.8 | 63 |
| LH2 B800-B850 | 0.5 | 3.5 | 1.7 ± 0.7 | 63 |
| LH2 B850-B850 | 0.011 | 0.045 | 0.018 ± 0.006 | 126 |
| LH1 B880-B880 | 0.012 | 0.020 | 0.016 ± 0.002 | 64 |

Note: *n* denotes the number of selected directed donor–acceptor pairs. Candidate pairs were restricted to pigments within the same LH1 or LH2 ring with calculated transfer times of < 20 ps. For each donor pigment, the acceptor yielding the shortest time constant was selected. For B800–B800, B850–B850, and B880–B880 transfer, pairs were counted separately for each donor pigment, allowing both transfer directions to be retained. When two pigments were each other's fastest-transfer partner, the pair contributed two entries, one for each direction. For B800–B850 transfer, only B800 pigments were treated as donors; therefore, only the B800→B850 direction was included.
